# Noribogaine effects on wakefulness and sleep

**DOI:** 10.1101/2023.07.26.550725

**Authors:** Juan Pedro Castro-Nin, Diego Serantes, Paola Rodriguez, Bruno Gonzalez, Ignacio Carrera, Pablo Torterolo, Joaquín González

## Abstract

Ibogaine is a potent atypical psychedelic that has gained considerable attention due to its antiaddictive and antidepressant properties in preclinical and clinical studies. Previous research from our group showed that ibogaine suppresses sleep and produces an altered wakefulness state which resembles natural REM sleep. However, after systemic administration, ibogaine is rapidly metabolized to noribogaine, which also shows antiaddictive effects and a distinct pharmacological profile, making this drug a promising therapeutic candidate. Therefore, whether the sleep/wake alterations depend on ibogaine or its principal metabolite noribogaine remains unknown. To answer this question, we conducted polysomnographic recordings in rats following the administration of pure noribogaine. Our results show that noribogaine promotes wakefulness while reducing slow-wave sleep and blocking REM sleep. Thus, like ibogaine, noribogaine significantly alters the sleep-wake architecture, highlighting the possible role of serotonin reuptake inhibition as a likely candidate underlying the wake-promoting and REM sleep-suppressing effects.

## Introduction

*Ibogaine* is a naturally occurring atypical psychedelic alkaloid found in the root bark of the African shrub *Tabernanthe iboga* (Lavaud and Massiot, 2017), used over centuries for medicinal and ritual purposes in equatorial Africa. Over the last decades, it attracted considerable scientific interest because of its long-lasting antiaddictive properties. These have been evidenced in preclinical research involving self-administration paradigms of opioids, cocaine, alcohol, and nicotine (Alper et al., 1999; Belgers et al., 2016; Cappendijk and Dzoljic, 1993; Glick et al., 2006; He et al., 2005; Rezvani et al., 1995), and in human reports, observational studies and open-label clinical trials involving several drugs of abuse (Brown and Alper, 2018; Mash et al., 2018; Noller et al., 2018; Schenberg et al., 2014).

Following its systemic administration in humans and rodents, ibogaine is rapidly metabolized to noribogaine (Baumann et al., 2001a, 2001b), which has a longer half-life than the parent drug and also exhibits anti-addictive properties (Glue et al., 2016; Mash et al., 2016). Importantly, the concentration of this active metabolite in rodents already reaches meaningful brain concentrations less than two hours following the initial intraperitoneal (i.p.) ibogaine injection, even as early as 20 minutes (Rodriguez et al., 2020). Notably, noribogaine’s pharmacological profile differs from its parent drug in several key aspects (Baumann et al., 2001a; Coleman et al., 2019; Glick et al., 2006; Jacobs et al., 2007; Layer et al., 1996; Mash et al., 1995).

Noribogaine shows a lower NMDA affinity than ibogaine, and is a significantly more potent serotonin reuptake inhibitor. Regarding the interaction to the 5HT2A receptor, noribogaine possesses even a weaker affinity than ibogaine. Additionally, noribogaine shows a higher affinity for mu and kappa opioid receptors and the norepinephrine transporter. Thus, the effects of ibogaine administration are mediated by an ibogaine/noribogaine combination in the brain and, as such, we still do not fully comprehend how each drug selectively affects brain function.

Moreover, while ibogaine’s effects involve an intense altered state of consciousness that resembles an oneiric experience (Brown et al., 2019; J. González et al., 2021; Naranjo, 2008), such psychedelic effects were not reported in a noribogaine clinical trial (Glue et al., 2016), although further studies using higher doses are needed to characterize the potential psychedelic properties of noribogaine. Recent evidence suggests that noribogaine might be the active antiaddictive agent mediating ibogaine’s neuroplastic effects (Marton et al., 2019), since only this drug (and not ibogaine) was capable of promoting structural neural plasticity in-vitro (Ly et al., 2018). Nevertheless, despite the comprehensive research performed on noribogaine action (Baumann et al., 2001a, 2001b; Coleman et al., 2019; Edinoff et al., 2022; Glue et al., 2016; Helsley et al., 1997; Iyer et al., 2021; Layer et al., 1996; Maillet et al., 2015; Mash et al., 1995; Moliner et al., 2023; Rodriguez et al., 2020), suggesting a promising antiaddictive profile, we still ignore how ibogaine’s principal metabolite affects sleep-wake patterns.

Hence, in this work, we performed polysomnographic recordings in chronically prepared rats in order to study the acute effects of noribogaine. We found that noribogaine enhanced wakefulness and decreased sleep. Moreover, similar to ibogaine, noribogaine persistently suppressed REM sleep. Thus, our results shed new light on the effects of noribogaine on the rat’s brain.

## Materials and Methods

### Experimental Animals

Nine Sprague Dawley male adult rats were maintained on a 12-h light/dark cycle (lights on at 06.00 AM). Food and water were freely available. The animals were determined to be in good health by veterinarians of the institution. All experimental procedures were conducted in agreement with the National Animal Care Law (No. 18611) and with the “Guide to the care and use of laboratory animals”(8th edition, National Academy Press, Washington DC, 2010).

Furthermore, the Institutional Animal Care Committee approved the experimental procedures (Exp.No. 070153-000332-1). Adequate measures were taken to minimize the pain, discomfort, or stress of the animals, and all efforts were made to use the minimal number of animals necessary to obtain reliable scientific data.

### Surgical Procedures

The animals were chronically implanted with electrodes to monitor the states of sleep and wakefulness. We employed similar surgical procedures as in our previous studies (González et al., 2018; J. González et al., 2021; Mondino et al., 2020). Anesthesia was induced with a mixture of ketamine-xylazine (110 mg/kg; 10 mg/kg i.p., respectively). The rat was positioned in a stereotaxic frame and the skull was exposed. To record the electroencephalogram, stainless steel screw electrodes were placed in the skull above the primary motor (M1) and somatosensory (S1) cortices, as well as over the cerebellum (reference electrode). To record electromyogram (EMG), a bipolar electrode was inserted into the neck muscle (Fig. 1A). The electrodes were soldered into a socket and fixed onto the skull with acrylic cement. At the end of the surgical procedures, they were adapted to the recording chamber for at least 4 days.

**Figure 1.**
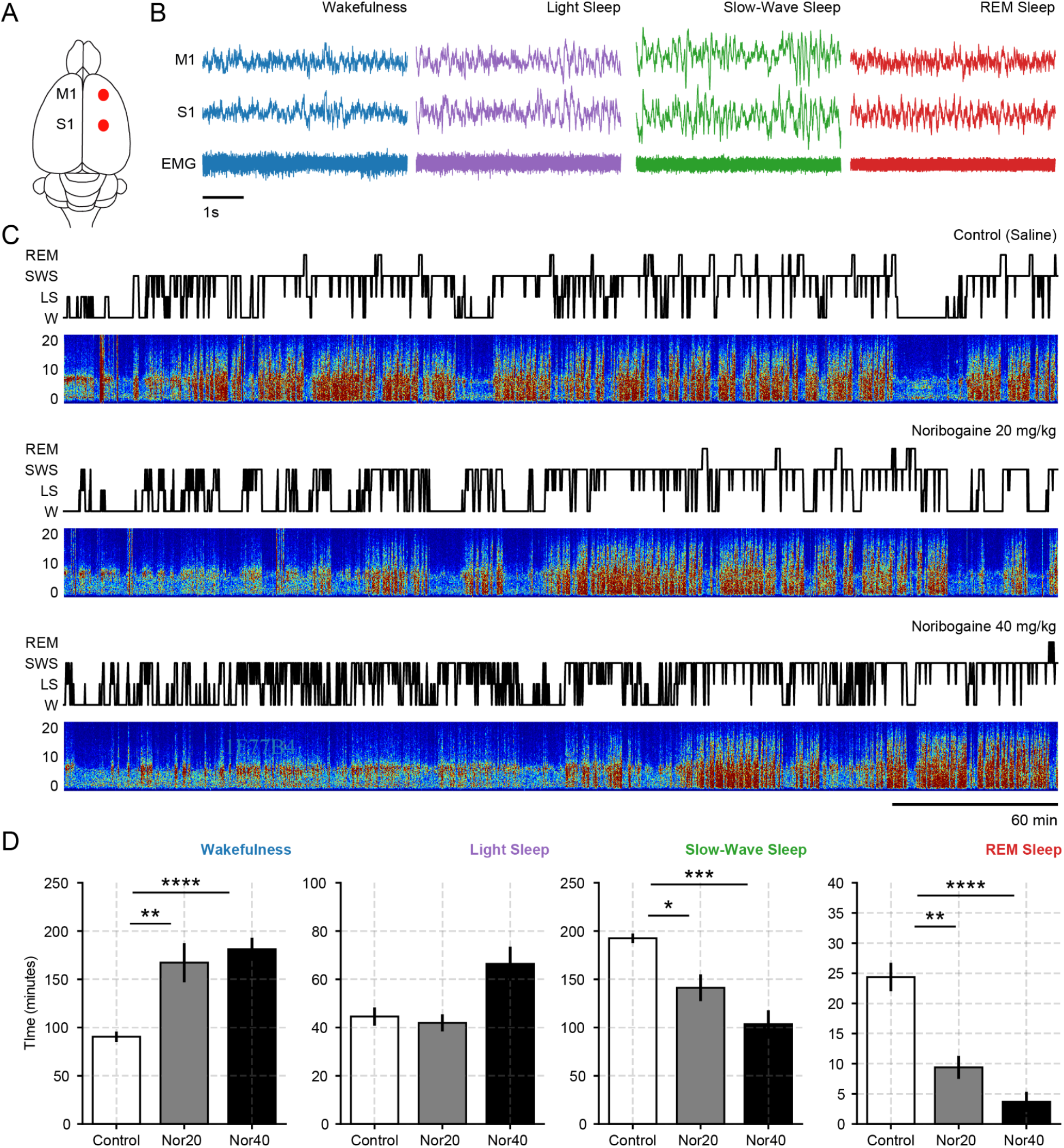
Noribogaine promotes wakefulness. **A** Location of the recording electrodes in the rat’s brain. M1: primary motor cortex; S1: primary somatosensory cortex; EMG: electromyogram. **B** Example polysomnographic recordings of each sleep-wake state obtained from a representative animal. A five-second window is shown, which clearly depicts the characteristic activity during each state. **C** Hypnograms and spectrograms from the same representative animal (panel B) during the whole six-hour recording session. Recordings started following saline (control), noribogaine 20 and 40 mg/kg i.p. administration. **D** Time spent in each sleep-wake state following control or noribogaine administration. The whole six-hour recordings were analyzed. Bar plots show the mean ± SEM. N: Control = 9; N_20_ = 6; N_40_ = 7. * p<0.05, ** p<0.01, *** p<0.001, **** p<0.00001, corrected for multiple comparisons through the Benjamini-Hochberg procedure.

**Figure 2.**
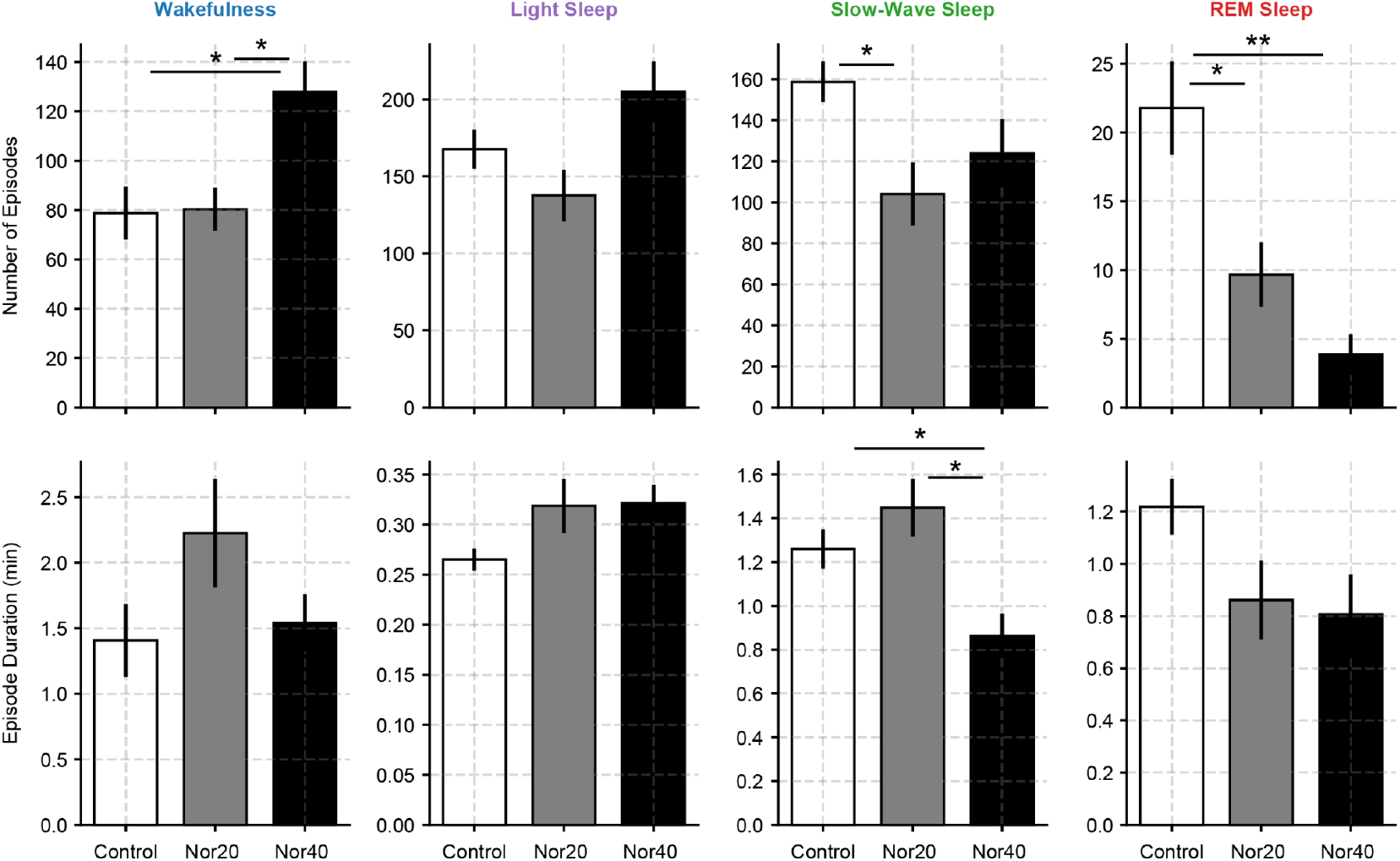
Noribogaine fragments Slow-Wave sleep. Top: Number of episodes of each sleep-wake state. Bottom: Average duration of the sleep-wake episodes. Episodes are defined by concatenating consecutive, non-interrupted, 10 s epochs of a single state. The whole six-hour recordings were analyzed. Bar plots show the mean ± SEM. N: Control = 9; N_20_ = 6; N_40_ = 7. * p<0.05, ** p<0.01, corrected for multiple comparisons through the Benjamini-Hochberg procedure.

### Experimental Sessions

Animals were housed individually in transparent cages (40 × 30 × 20 cm) containing wood shaving material in a temperature-controlled room (21-24 °C), with water and food *ad libitum*. Experimental sessions were conducted during the light period, between 11 AM and 5 PM in a sound-attenuated chamber with Faraday shield. Before the beginning of the recordings, animals were injected with a noribogaine dose or vehicle i.p. The recordings were performed through a rotating commutator, to allow the rats to freely move within the recording box.

Polysomnographic data was amplified (x 1000), sampled at 1024 Hz with a 16-bit AD converter, and stored in a PC using DasyLab Software.

### Noribogaine administration

Noribogaine·HCl was prepared in the Laboratorio de Síntesis Orgánica-Facultad de Química, Universidad de la República, by O-demethylation of ibogaine (obtained by decarboxylation of voacangine isolated from *Voacanga africana* root bark according to our previously reported method) (B. González et al., 2021). In brief, in a two neck round bottom flask under an argon atmosphere, ibogaine (493 mg, 1.58 mmol) was dissolved in dry 1,2-dichloroethane (15.8 mL, 0.1 M) using magnetic stirring. Then, ethanetiol (497 μL, 6.34 mmol, 4.0 eq) was added followed by the addition of a 1.0 M solution of boron tribromide (2,38 mL, 1.5eq). The system was heated to 55°C for 2 hours, when total consumption of the starting material was evidenced by thin layer chromatography (TLC). The reaction was quenched by adding methanol until all the suspended solids were dissolved. The resulting reaction mixture was diluted with EtOAc, and transferred to a separation funnel where a saturated sodium bicarbonate solution was added. The aqueous layer was extracted with ethyl acetate (EtOAc x3), and the combined organic layers were dried over sodium sulfate. The solvent was removed under vacuum to obtain a crude mixture which was purified by column chromatography [SiO_2_; 50% EtOAc in Hexane, 1% NH_4_OH_(cc)_].

Noribogaine free base was obtained as a white amorphous solid (420 mg, 89% yield). To prepare the corresponding hydrochloride, the free base was dissolved in dry diethyl ether and an anhydrous solution of HCl in diethyl ether (3 M, 0.75 mL, 1.5eq) was added. A white solid was formed, which was filtrated and washed several times with dry diethyl ether. The resulting Noribogaine-HCl (425 mg) was dried under vacuum and obtained as a white solid.

Dissolution of noribogaine·HCl to prepare the samples for i.p. injection was carried out using a warm saline vehicle that was previously degassed by nitrogen bubbling. In order to study the effects of noribogaine on sleep and wakefulness, we employed an unpaired design, in which at the beginning of the recordings, rats either received noribogaine 20 mg/kg (N_20_, six rats), noribogaine 40 mg/kg (N_40_, seven rats), or saline solution (Control, nine rats) i.p.. These doses have been used in previous preclinical studies (Baumann et al., 2001b; Glick et al., 2006; Rodriguez et al., 2020). Out of all the animals, four received both doses on different days in a counterbalanced order, while two extra animals received N_20_ and three extra N_40_. The wash-out period between doses was three days.

### Sleep Scoring

The states of sleep and wakefulness were determined in 10-s epochs. Wakefulness (W) was defined as low-voltage fast waves in the motor cortex, a noticeable theta rhythm (4-7 Hz) in S1 and relatively high EMG activity. Light Sleep (LS) was defined by high amplitude slow cortical waves intermixed by low amplitude fast EEG activity. Slow-Wave Sleep (SWS) was determined by the presence of high-voltage slow cortical waves together with sleep spindles in M1 and S1 associated with a reduced EMG amplitude; REM sleep as low voltage fast frontal waves, a regular theta rhythm in S1 and a silent EMG except for occasional twitches (Fig. 1B). Total time spent in W, LS, SWS, and REM sleep, as well as the duration and number of episodes over a 6 h recording period was analyzed. Sleep latencies were also evaluated.

### Statistics

All values are presented as mean ± SEM. The experimental design for the sleep analysis was a within-subject design, where the statistical significance of the differences among groups (noribogaine 0, 20, 40 mg/kg) was evaluated utilizing an unpaired one-way analysis of variance (ANOVA) and a student t-test as a *post hoc* test. The resulting p values were corrected for multiple comparisons through the Benjamini-Hochberg False Discovery Rates procedure.

Statistical significance was set at p<0.05. We reported the full ANOVA result and the corresponding significant p-values for the *post hoc* test.

## Results

### Noribogaine promotes wakefulness and impairs sleep

Figure 1C shows the hypnograms and spectrograms of a representative animal following control, N_20_ and N_40_ conditions. This representative example shows that both noribogaine doses drastically altered the sleep-wake patterns, favoring wakefulness and decreasing sleep. Group results during the whole six-hour recordings confirmed that both noribogaine doses significantly promoted W (F(2,19) = 14.01, p = 0.0001, Control vs. N_20_, p = 0.002; Control vs. N_40_, p = 0.00005) while significantly reducing SWS (F(2,19) = 11.99, p = 0.0004, Control vs. N_20_, p = 0.01; Control vs. N_40_, p = 0.0002) and REM (F(2,19) = 21.34, p = 0.00001, Control vs. N_20_, p = 0.002; Control vs. N_40_, p = 0.00007).

Next, having observed a pronounced change in the sleep architecture, we analyzed the number and duration of episodes for each sleep-wake state. *Episodes* are defined as consecutive, non-interrupted single-state epochs. We found that noribogaine increased the number of W episodes (F(2,19) = 5.51, p = 0.012, Control vs. N_40_, p = 0.024; N_20_ vs. N_40_, p = 0.024) and reduced SWS and REM episodes (SWS: F(2,19) = 3.67, p = 0.044, Control vs. N_20_, p = 0.036; REM: F(2,19) = 10.34, p = 0.0009, Control vs. N_20_, p = 0.042; Control vs. N_40_, p = 0.003). In contrast, the average duration of each episode remained similar, with the exception of the N_40_ dose, which significantly reduced the duration of the SWS episodes (F(2,19) = 6.39, p = 0.007, Control vs. N_40_, p = 0.023; N_20_ vs. N_40_, p = 0.021).

### Noribogaine effects peak during the first two hours

After analyzing the whole recording, we sought to disentangle the time-dependent effects of noribogaine action. Hence, we divided our recordings into three blocks of 2 hours. When considering the effects on each 2-hour block, we found that the peak effects for both doses occurred within the first 2 hours following the i.p. administration (Fig 3). Specifically, both doses promoted W (F(2,19) = 18.44, p = 0.00003, Control vs. N_20_, p = 0.0001; Control vs. N_40_, p = 0.0001) while decreasing SWS (F(2,19) = 25.07, p = 0.000004, Control vs. N_20_, p = 0.00002; Control vs. N_40_, p = 0.00004) and REM (F(2,19) = 8.73, p = 0.002, Control vs. N_20_, p = 0.020; Control vs. N_40_, p = 0.020).

**Figure 3.**
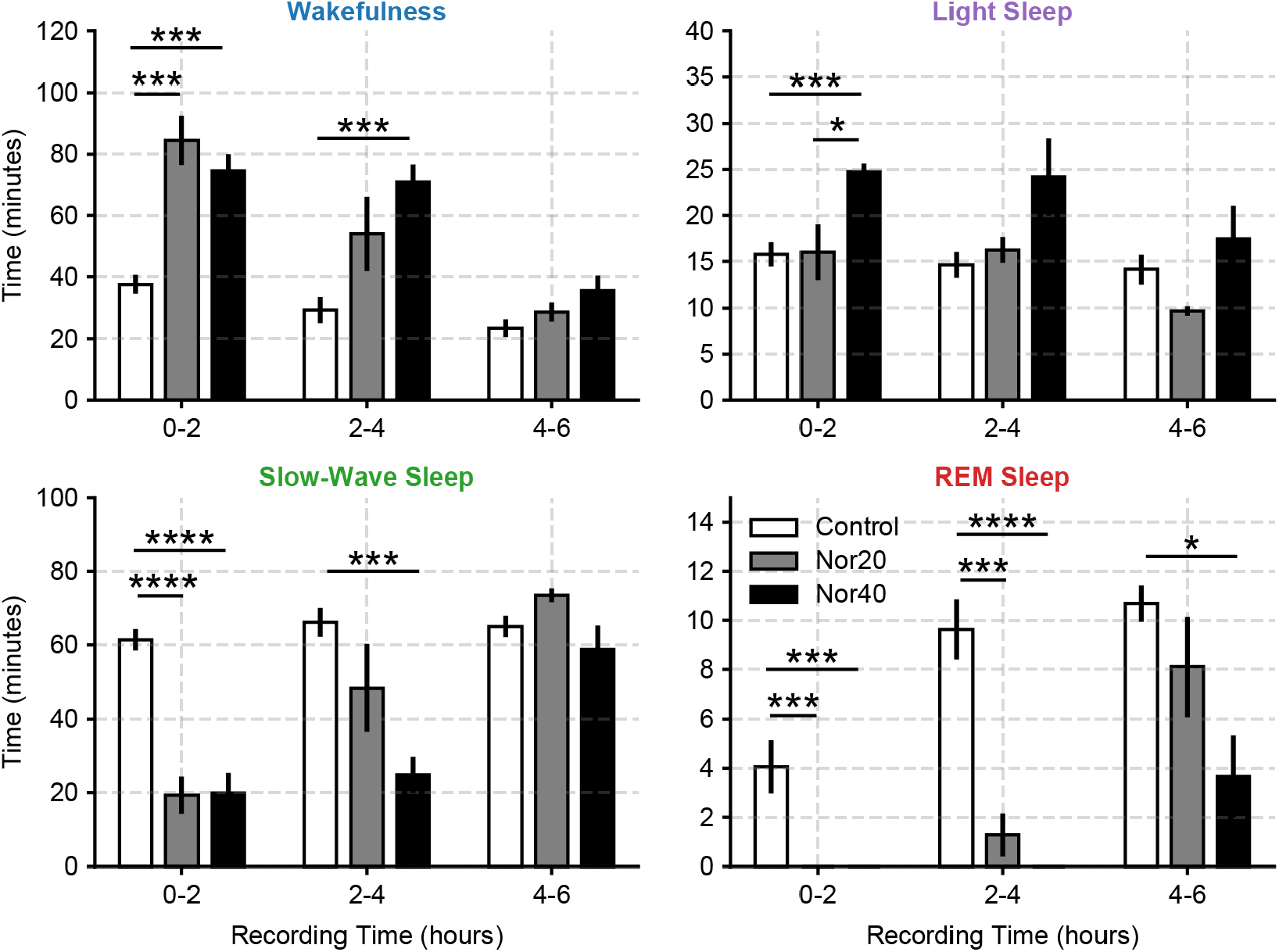
Noribogaine produces a long-term REM sleep suppression. Time spent in each sleep-wake state following control or noribogaine administration. The whole six-hour recording was divided in two hour blocks. Bar plots show the mean ± SEM. N: Control = 9; N_20_ = 6; N_40_ = 7. * p<0.05, ** p<0.01, *** p<0.001, **** p<0.00001, corrected for multiple comparisons through the Benjamini-Hochberg procedure.

This profile was conserved for the higher dose during the 2-4 hour block, while we still observed a significant REM reduction for the 20 mg dose (W: F(2,19) = 7.59, p = 0.0037, Control vs. N_40_, p = 0.0003. SWS: F(2,19) = 8.18, p = 0.0027, Control vs. N_40_, p = 0.0001. REM: F(2,19) = 28.10, p = 0.000002, Control vs. N_20_, p = 0.0006; Control vs. N_40_, p = 0.00004). In contrast, the last two-hour block only showed a significant reduction of REM sleep for the larger noribogaine dose (F(2,19) = 5.04, p = 0.017, Control vs. N_40_, p = 0.010), further evidencing the dose dependency of noribogaine effects. Therefore, our results suggest that noribogaine effects peak during the first 2 hours and are slowly washed-out in the later hours.

### Noribogaine suppresses REM sleep initiation

Finally, motivated by the long-lasting REM suppression, we studied the latency of REM initiation. First, Figure 4A shows a representative hypnogram from an animal treated with control and both noribogaine doses. We can observe that the first control REM episode occurs 80 minutes after saline injection. Surprisingly, the first REM episode occurred after 5 hours in the 20 mg dose and did not even occur within the six-hour recording for the larger 40 mg dose.

**Figure 4.**
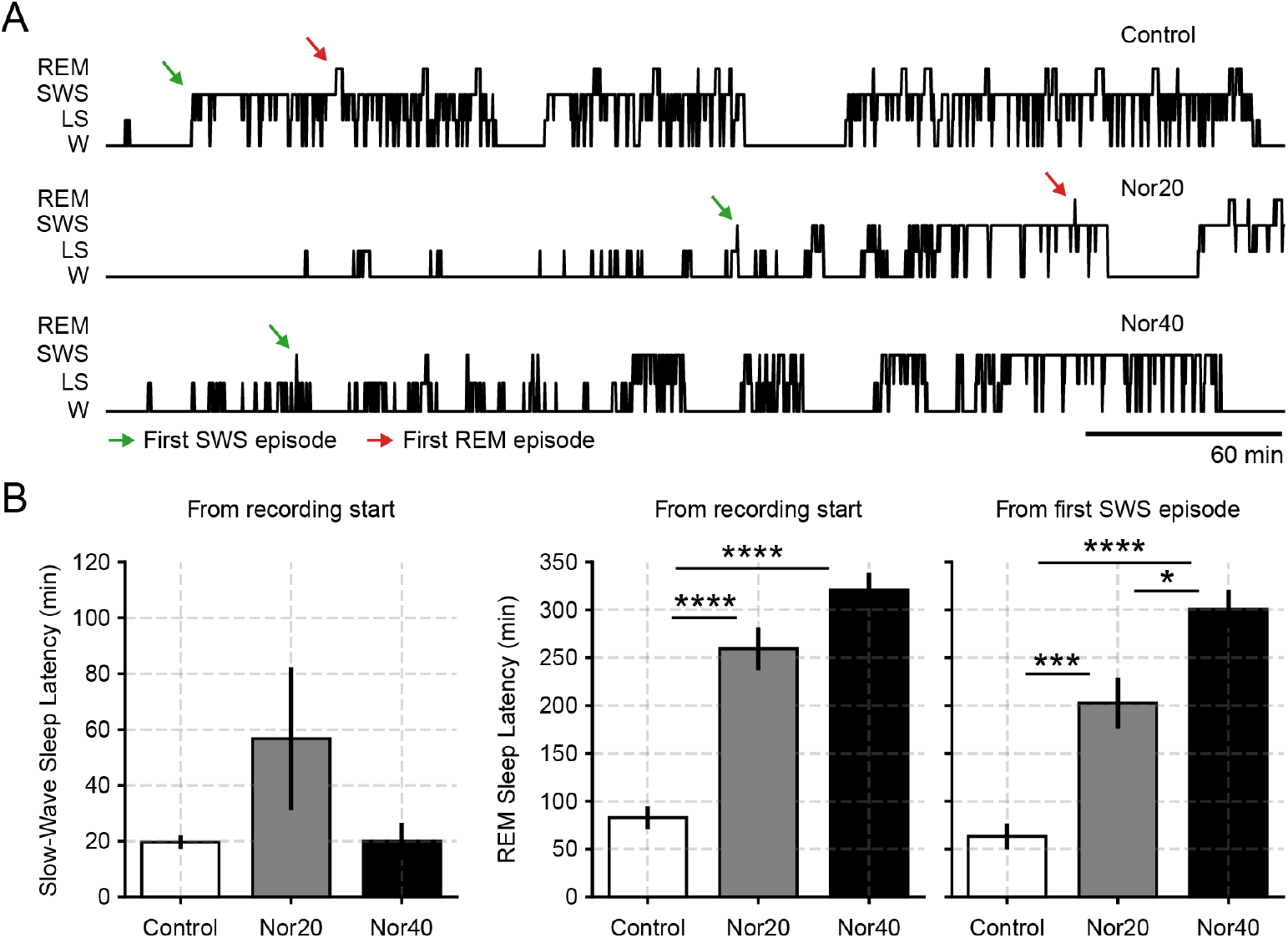
Noribogaine delays REM sleep initiation. **A** Hypnograms of a representative animal are shown after saline, noribogaine 20, and noribogaine 40 mg/kg i.p. Administration. The whole six-hour recordings are shown. The arrowheads above the hypnogram show either the SWS (green) or REM (red) latencies. Notice that the 40 mg dose completely suppressed REM in this example. **B** Latencies to SWS (left), REM (middle), and to REM from the first SWS epoch (right) are shown. Bar plots show the mean ± SEM. N: Control = 9; N_20_ = 6; N_40_ = 7. * p<0.05, ** p<0.01, *** p<0.001, **** p<0.00001, corrected for multiple comparisons through the Benjamini-Hochberg procedure.

Group data confirmed this trend, where we found that REM latency significantly increased for both doses (Fig 4B, left panel, F(2,19) = 50.18, p = 0.00000002, Control vs. N_20_, p = 0.00001; Control vs. N_40_, p = 0.0000001). Moreover, when we computed the REM latency from the first SWS episode, we found a dose-dependent pattern in which the latency increased from control to N_20_ and N_20_ to N_40_ (Fig 4B, right panel, F(2,19) = 35.80, p = 0.0000003, Control vs. N_20_, p = 0.0004; Control vs. N_40_, p = 0.0000005, N_20_ vs. N_40_, p = 0.018). Noteworthy, noribogaine did not significantly alter the SWS latency, confirming the specificity of the effects on REM.

## Discussion

In this work, we found that noribogaine increased wakefulness to the detriment of sleep. Importantly, while the SWS effects occurred close to the noribogaine injection, REM was persistently blocked throughout the recording session. Thus, our results confirm that similar to ibogaine (González et al., 2018), noribogaine also profoundly perturbs the sleep-wake architecture.

When comparing our results to our previous ibogaine experiments, we note that the larger noribogaine dose – commonly employed in addiction protocols (Baumann et al., 2001b; Glick et al., 2006; Rodriguez et al., 2020) – elicited a higher concentration peak in brain tissue than the concentration observed after ibogaine administration (Rodriguez et al., 2020). However, despite the pharmacological differences between ibogaine and its principal metabolite, their sleep-wake effects shared two key aspects: 1) both drugs promoted W while impairing normal SWS. These effects were more noticeable during the first two hours and gradually disappeared during the course of the recording period. 2) both drugs significantly delayed REM onset. However, while W-promoting effects were comparable after both drugs administration, noribogaine’s effects on REM were appreciably larger than ibogaine’s.

These results suggest that the serotonin transporter (SERT) blockade likely mediates the W-promoting and REM-suppressing effects of both drugs. We reasoned that if serotonergic effects were to mediate both alterations, noribogaine should show larger effects due to its more potent serotonin reuptake inhibition (Coleman et al., 2019; Jacobs et al., 2007). In support of this idea, the significant W-promoting effects mediated by noribogaine were significant up to the second recording block, while ibogaine’s effects were restricted to the first 2 hours period only (see Gonzalez et al., 2018. Figure 2). Moreover, both doses of noribogaine completely suppressed REM during the first 2-hour block, while REM did occur during this first period under ibogaine, albeit in a reduced proportion (Fig. 2, González et al., 2018). Furthermore, while total REM time reached almost 11 minutes during the entire recording for the larger ibogaine dose (Table. 1, González et al., 2018), it did not even reach 5 minutes during the equivalent noribogaine dose. Hence, by comparing noribogaine results with ibogaine, we argue that common mechanisms mediate these drugs’ W-promoting and REM-suppressing effects.

Nevertheless, detailed studies comparing equal brain concentrations of both compounds are needed to elucidate which common/independent mechanisms underpin their sleep-wake alterations.

We cannot discard that several neuromodulators may also contribute to the observed sleep-wake alterations. For instance, cholinergic effects should also be considered as a common activating pathway for both ibogaine and noribogaine (Da Costa et al., 1980; Da Costa-Rochette et al., 1981; Schneider and Sigg, 1957), since mesopontine and basal forebrain cholinergic neurons are involved in the generation and maintenance of W (Torterolo et al., 2016). Furthermore, opioid receptors effects should also be considered since opioid abuse alters sleep-wake architecture (Hartwell et al., 2014; Robertson et al., 2016). Nevertheless, due to ibogaine’s and noribogaine’s complex pharmacology, the interactions between these and other neurotransmitters should be considered when interpreting these results.

In recent years, there has been an effort to improve psychedelics by removing some of their undesired side effects (i.e., toxicity, perceptual alterations) while keeping their positive health effects (Cameron et al., 2021). These compounds’ potential health benefits include treating depression, anxiety, and addictions, outperforming traditional therapeutics (Palhano-Fontes et al., 2022, 2019). These benefits depend on long-lasting neuroplastic processes elicited by these promising drugs (Cameron et al., 2021; Vargas et al., 2023). Of interest, noribogaine does not produce the behavioral effects that ibogaine produces, such as the flat body posture or tremors, while keeping its antidepressive and antiaddictive properties (Rodriguez et al., 2020). Moreover, It is important to note that at lower doses, ibogaine typically displays stimulant effects, while these were not reported for noribogaine (Glue et al., 2016). However, further studies employing higher doses are needed to clarify noribogaine’s potential psychedelic effects since ibogaine’s psychedelic effects are evidenced at higher drug concentrations. Still, our work shows that sleep-wake alterations are also present for noribogaine. Notably, traditional serotonergic antidepressants also blockade REM, confirming the ties between noribogaine’s pharmacological targets and its antidepressive profile (Rodriguez et al., 2020). Thus, sleep-wake alterations must be considered when employing noribogaine to treat substance abuse disorders.

Overall, our results demonstrate that noribogaine shares many similarities with its parent drug, suggesting that common mechanisms at different intensities are involved in the behavioral effects of both drugs. Hence, sleep perturbations should be considered when employing this drug in further clinical trials as a potential treatment for substance abuse disorder.

## Notes

### Competing Interest Statement

The authors have declared no competing interest.

## References

Alper, K.R., Lotsof, H.S., Frenken, G.M., Luciano, D.J., Bastiaans, J., 1999. Treatment of acute opioid withdrawal with ibogaine. Am. J. Addict. 8, 234–242.

Baumann, M.H., Pablo, J., Ali, S.F., Rothman, R.B., Mash, D.C., 2001a. Comparative neuropharmacology of ibogaine and its O-desmethyl metabolite, noribogaine. Alkaloids Chem. Biol. 56, 79–113.

Baumann, M.H., Rothman, R.B., Pablo, J.P., Mash, D.C., 2001b. In vivo neurobiological effects of ibogaine and its O-desmethyl metabolite, 12-hydroxyibogamine (noribogaine), in rats. J. Pharmacol. Exp. Ther. 297, 531–539.

Belgers, M., Leenaars, M., Homberg, J.R., Ritskes-Hoitinga, M., Schellekens, A.F.A., Hooijmans, C.R., 2016. Ibogaine and addiction in the animal model, a systematic review and meta-analysis. Transl. Psychiatry 6, e826.

Brown, T.K., Alper, K., 2018. Treatment of opioid use disorder with ibogaine: detoxification and drug use outcomes. Am. J. Drug Alcohol Abuse 44, 24–36.

Brown, T.K., Noller, G.E., Denenberg, J.O., 2019. Ibogaine and Subjective Experience: Transformative States and Psychopharmacotherapy in the Treatment of Opioid Use Disorder. J. Psychoactive Drugs 51, 155–165.

Cameron, L.P., Tombari, R.J., Lu, J., Pell, A.J., Hurley, Z.Q., Ehinger, Y., Vargas, M.V., McCarroll, M.N., Taylor, J.C., Myers-Turnbull, D., Liu, T., Yaghoobi, B., Laskowski, L.J., Anderson, E.I., Zhang, G., Viswanathan, J., Brown, B.M., Tjia, M., Dunlap, L.E., Rabow, Z.T., Fiehn, O., Wulff, H., McCorvy, J.D., Lein, P.J., Kokel, D., Ron, D., Peters, J., Zuo, Y., Olson, D.E., 2021. A non-hallucinogenic psychedelic analogue with therapeutic potential. Nature 589, 474–479.

Cappendijk, S.L., Dzoljic, M.R., 1993. Inhibitory effects of ibogaine on cocaine self-administration in rats. Eur. J. Pharmacol. 241, 261–265.

Coleman, J.A., Yang, D., Zhao, Z., Wen, P.-C., Yoshioka, C., Tajkhorshid, E., Gouaux, E., 2019. Serotonin transporter-ibogaine complexes illuminate mechanisms of inhibition and transport. Nature 569, 141–145.

Da Costa, L., Sulklaper, I., Naquet, R., 1980. [Modification of awake-sleep equilibrium by tabernanthine and some of its derivatives in the cat (author’s transl)]. Rev. Electroencephalogr. Neurophysiol. Clin. 10, 105–112.

Da Costa-Rochette, L., Sulklaper, I., Tomei, C., Naquet, R., 1981. [Restoration of sleep in cats pretreated with tabernanthine p-chlorophenoxyacetate (SAD 103) (author’s transl)]. Rev. Electroencephalogr. Neurophysiol. Clin. 11, 147–154.

Edinoff, A.N., Wu, N.W., Nix, C.A., Bonin, B., Mouhaffel, R., Vining, S., Gibson, W., Cornett, E.M., Murnane, K.S., Kaye, A.M., Kaye, A.D., 2022. Historical Pathways for Opioid Addiction, Withdrawal with Traditional and Alternative Treatment Options with Ketamine, Cannabinoids, and Noribogaine: A Narrative Review. Health Psychol Res 10, 38672.

Glick, S.D., Maisonneuve, I.M., Hough, L.B., Kuehne, M.E., Bandarage, U.K., 2006. (±)-18-methoxycoronaridine: A novel iboga alkaloid congener having potential anti-addictive efficacy. CNS Drug Rev. 5, 27–42.

Glue, P., Cape, G., Tunnicliff, D., Lockhart, M., Lam, F., Hung, N., Hung, C.T., Harland, S., Devane, J., Crockett, R.S., Howes, J., Darpo, B., Zhou, M., Weis, H., Friedhoff, L., 2016. Ascending Single-Dose, Double-Blind, Placebo-Controlled Safety Study of Noribogaine in Opioid-Dependent Patients. Clin Pharmacol Drug Dev 5, 460–468.

González, B., Fagúndez, C., Peixoto de Abreu Lima, A., Suescun, L., Sellanes, D., Seoane, G.A., Carrera, I., 2021. Efficient Access to the Iboga Skeleton: Optimized Procedure to Obtain Voacangine from Root Bark. ACS Omega 6, 16755–16762.

González, J., Cavelli, M., Castro-Zaballa, S., Mondino, A., Tort, A.B.L., Rubido, N., Carrera, I., Torterolo, P., 2021. EEG Gamma Band Alterations and REM-like Traits Underpin the Acute Effect of the Atypical Psychedelic Ibogaine in the Rat. ACS Pharmacol Transl Sci 4, 517–525.

González, J., Prieto, J.P., Rodríguez, P., Cavelli, M., Benedetto, L., Mondino, A., Pazos, M., Seoane, G., Carrera, I., Scorza, C., Torterolo, P., 2018. Ibogaine Acute Administration in Rats Promotes Wakefulness, Long-Lasting REM Sleep Suppression, and a Distinctive Motor Profile. Front. Pharmacol. 9, 374.

Hartwell, E.E., Pfeifer, J.G., McCauley, J.L., Moran-Santa Maria, M., Back, S.E., 2014. Sleep disturbances and pain among individuals with prescription opioid dependence. Addict. Behav. 39, 1537–1542.

He, D.-Y., McGough, N.N.H., Ravindranathan, A., Jeanblanc, J., Logrip, M.L., Phamluong, K., Janak, P.H., Ron, D., 2005. Glial cell line-derived neurotrophic factor mediates the desirable actions of the anti-addiction drug ibogaine against alcohol consumption. J. Neurosci. 25, 619–628.

Helsley, S., Rabin, R.A., Winter, J.C., 1997. The effects of noribogaine and harmaline in rats trained with ibogaine as a discriminative stimulus. Life Sci. 60, PL147–53.

Iyer, R.N., Favela, D., Zhang, G., Olson, D.E., 2021. The iboga enigma: the chemistry and neuropharmacology of iboga alkaloids and related analogs. Nat. Prod. Rep. 38, 307–329.

Jacobs, M.T., Zhang, Y.-W., Campbell, S.D., Rudnick, G., 2007. Ibogaine, a noncompetitive inhibitor of serotonin transport, acts by stabilizing the cytoplasm-facing state of the transporter. J. Biol. Chem. 282, 29441–29447.

Lavaud, C., Massiot, G., 2017. The Iboga Alkaloids. Prog. Chem. Org. Nat. Prod. 105, 89–136.

Layer, R.T., Skolnick, P., Bertha, C.M., Bandarage, U.K., Kuehne, M.E., Popik, P., 1996. Structurally modified ibogaine analogs exhibit differing affinities for NMDA receptors. Eur. J. Pharmacol. 309, 159–165.

Ly, C., Greb, A.C., Cameron, L.P., Wong, J.M., Barragan, E.V., Wilson, P.C., Burbach, K.F., Soltanzadeh Zarandi, S., Sood, A., Paddy, M.R., Duim, W.C., Dennis, M.Y., McAllister, A.K., Ori-McKenney, K.M., Gray, J.A., Olson, D.E., 2018. Psychedelics Promote Structural and Functional Neural Plasticity. Cell Rep. 23, 3170–3182.

Maillet, E.L., Milon, N., Heghinian, M.D., Fishback, J., Schürer, S.C., Garamszegi, N., Mash, D.C., 2015. Noribogaine is a G-protein biased κ-opioid receptor agonist. Neuropharmacology 99, 675–688.

Marton, S., González, B., Rodríguez-Bottero, S., Miquel, E., Martínez-Palma, L., Pazos, M., Prieto, J.P., Rodríguez, P., Sames, D., Seoane, G., Scorza, C., Cassina, P., Carrera, I., 2019. Ibogaine Administration Modifies GDNF and BDNF Expression in Brain Regions Involved in Mesocorticolimbic and Nigral Dopaminergic Circuits. Front. Pharmacol. 10, 193.

Mash, D.C., Ameer, B., Prou, D., Howes, J.F., Maillet, E.L., 2016. Oral noribogaine shows high brain uptake and anti-withdrawal effects not associated with place preference in rodents. J. Psychopharmacol. 30, 688–697.

Mash, D.C., Duque, L., Page, B., Allen-Ferdinand, K., 2018. Ibogaine Detoxification Transitions Opioid and Cocaine Abusers Between Dependence and Abstinence: Clinical Observations and Treatment Outcomes. Front. Pharmacol. 9, 529.

Mash, D.C., Staley, J.K., Pablo, J.P., Holohean, A.M., Hackman, J.C., Davidoff, R.A., 1995. Properties of ibogaine and its principal metabolite (12-hydroxyibogamine) at the MK-801 binding site of the NMDA receptor complex. Neurosci. Lett. 192, 53–56.

Moliner, R., Girych, M., Brunello, C.A., Kovaleva, V., Biojone, C., Enkavi, G., Antenucci, L., Kot, E.F., Goncharuk, S.A., Kaurinkoski, K., Kuutti, M., Fred, S.M., Elsilä, L.V., Sakson, S., Cannarozzo, C., Diniz, C.R.A.F., Seiffert, N., Rubiolo, A., Haapaniemi, H., Meshi, E., Nagaeva, E., Öhman, T., Róg, T., Kankuri, E., Vilar, M., Varjosalo, M., Korpi, E.R., Permi, P., Mineev, K.S., Saarma, M., Vattulainen, I., Casarotto, P.C., Castrén, E., 2023. Psychedelics promote plasticity by directly binding to BDNF receptor TrkB. Nat. Neurosci. 26, 1032–1041.

Mondino, A., Cavelli, M., González, J., Osorio, L., Castro-Zaballa, S., Costa, A., Vanini, G., Torterolo, P., 2020. Power and Coherence in the EEG of the Rat: Impact of Behavioral States, Cortical Area, Lateralization and Light/Dark Phases. Clocks Sleep 2, 536–556.

Naranjo, C., 2008. Psycotherapeutic Possibilities of New Fantasy-Enhancing Drugs. Clin. Toxicol. https://doi.org/10.3109/15563656908990930

Noller, G.E., Frampton, C.M., Yazar-Klosinski, B., 2018. Ibogaine treatment outcomes for opioid dependence from a twelve-month follow-up observational study. Am. J. Drug Alcohol Abuse 44, 37–46.

Palhano-Fontes, F., Barreto, D., Onias, H., Andrade, K.C., Novaes, M.M., Pessoa, J.A., Mota-Rolim, S.A., Osório, F.L., Sanches, R., Dos Santos, R.G., Tófoli, L.F., de Oliveira Silveira, G., Yonamine, M., Riba, J., Santos, F.R., Silva-Junior, A.A., Alchieri, J.C., Galvão-Coelho, N.L., Lobão-Soares, B., Hallak, J.E.C., Arcoverde, E., Maia-de-Oliveira, J.P., Araújo, D.B., 2019. Rapid antidepressant effects of the psychedelic ayahuasca in treatment-resistant depression: a randomized placebo-controlled trial. Psychol. Med. 49, 655–663.

Palhano-Fontes, F., Soares, B.L., Galvão-Coelho, N.L., Arcoverde, E., Araujo, D.B., 2022. Ayahuasca for the Treatment of Depression. Curr. Top. Behav. Neurosci. 56, 113–124.

Rezvani, A.H., Overstreet, D.H., Lee, Y.W., 1995. Attenuation of alcohol intake by ibogaine in three strains of alcohol-preferring rats. Pharmacol. Biochem. Behav. 52, 615–620.

Robertson, J.A., Purple, R.J., Cole, P., Zaiwalla, Z., Wulff, K., Pattinson, K.T.S., 2016. Sleep disturbance in patients taking opioid medication for chronic back pain. Anaesthesia 71, 1296–1307.

Rodriguez, P., Urbanavicius, J., Prieto, J.P., Fabius, S., Reyes, A.L., Havel, V., Sames, D., Scorza, C., Carrera, I., 2020. A Single Administration of the Atypical Psychedelic Ibogaine or Its Metabolite Noribogaine Induces an Antidepressant-Like Effect in Rats. ACS Chem. Neurosci. 11, 1661–1672.

Schenberg, E.E., de Castro Comis, M.A., Chaves, B.R., da Silveira, D.X., 2014. Treating drug dependence with the aid of ibogaine: a retrospective study. J. Psychopharmacol. 28, 993–1000.

Schneider, J.A., Sigg, E.B., 1957. Neuropharmacological studies on ibogaine, an indole alkaloid with central-stimulant properties. Ann. N. Y. Acad. Sci. 66, 765–776.

Torterolo, P., Monti, J.M., Pandi-Perumal, S.R., 2016. Chapter 1 neuroanatomy and neuropharmacology of sleep and wakefulness, in: Synopsis of Sleep Medicine. Apple Academic Press, pp. 1–22.

Vargas, M.V., Dunlap, L.E., Dong, C., Carter, S.J., Tombari, R.J., Jami, S.A., Cameron, L.P., Patel, S.D., Hennessey, J.J., Saeger, H.N., McCorvy, J.D., Gray, J.A., Tian, L., Olson, D.E., 2023. Psychedelics promote neuroplasticity through the activation of intracellular 5-HT2A receptors. Science 379, 700–706.

